# Tamoxifen reduces inflammatory infiltration of neutrophils in the airways

**DOI:** 10.1101/2020.06.19.161919

**Authors:** Agustín Mansilla, Jaime Soto, Claudio Henríquez, Amber Philp, Marcus A. Mall, José Sarmiento, Carlos A. Flores

**Author notes:** Corresponding author: Carlos A. Flores, Centro de Estudios Científicos (CECs)., Valdivia, 511046, Chile.

## Abstract

Tamoxifen is a drug of choice for treatment of breast cancer but it has also been reported to bear anti-inflammatory activity. Previously, we have observed that tamoxifen treatment successfully reduced inflammation in horses affected by equine asthma syndrome. More notorious, tamoxifen was effective at reducing the infiltration of neutrophils in the inflamed airways by a mechanism that increases efferocytosis, allowing the resolution of inflammation. Due to the emerging increase in patients with chronic inflammatory lung diseases, there is an urgent need for therapies that help reduce airway inflammation. Thus we tested the effect of tamoxifen on airway neutrophil infiltration by using three different mouse models for acute and chronic inflammation of the lung. We found that the drug was able to produce a significant reduction in neutrophils in all scenarios. Ussing chamber experiments demonstrated that tamoxifen has no effect on fluid secretion and absorption, discarding a possible reduction in mucociliary clearance due to the known ion-channel blocking effects of tamoxifen. Since this drug has been largely used in human medicine, tamoxifen might be a “low hanging fruit” for a novel anti-inflammatory therapy for airway diseases characterized by neutrophilic inflammation.

## INTRODUCTION

Neutrophils are immune cells whose function is absolutely necessary for an efficient defensive response, nevertheless, an increase in neutrophil recruitment or a failure in their clearance is associated with collateral tissue damage, delayed or defective healing, and non-resolved inflammation [1,2]. For example, chronic inflammation as observed in cystic fibrosis (CF), non-eosinophilic asthma or chronic obstructive pulmonary disease (COPD), as well as acute inflammatory responses including Covid-19 are often affected by massive neutrophilic infiltration, thus its contention appears necessary [3–6]. In fact, animal models of acute lung injury (ALI) have provided insight as neutrophil depletion, or inhibition of neutrophil function, invariably ameliorates the extent of damage in the lungs [7]. In humans with neutrophilic asthma, many of the new therapeutic candidates, targets neutrophil’s arrival, correlating a lower neutrophilic infiltration of the airway with reduce exacerbations, improving lung function and quality of life [8].

We have previously determined that tamoxifen (TAMX) was effective in the management of horses affected by equine asthma syndrome (EAS), a naturally occurring asthma-like condition characterized by severe airway inflammation and neutrophilic infiltration. Clinical and endoscopic scores improved, along with decreased neutrophils levels in the broncoalveolar lavage fluid (BALF) in the animals that received the TAMX [9]. TAMX is a drug of choice in the treatment of breast cancer but also considered a nonsteroidal anti-inflammatory agent (NSAID) [10]. In this work, we tested the potential of TAMX to reduce neutrophil infiltration of the airways using different mouse models of airway inflammation including acute lung injury (ALI) and the *Scnnb1*^tg/+^ mouse model of COPD/CF, and tested TAMX properties as a blocker of ion channels in mouse tracheas using electrophysiology.

## METHODS

### Reagents

All reagents were obtained from Sigma-Aldrich unless stated otherwise.

### Animals

Mice were housed at CECs animal facility. The *Scnn1b*^tg/+^ mouse was backcrossed to the C57BL/6J background as previously described [11]. Animals were maintained in the Specific Pathogen Free mouse facility of Centro de Estudios Cientificos (CECs) with access to food and water *ad libitum*. 6-week-old male or female mice were used (C57BL/6J). All experimental procedures were approved by the Centro de Estudios Científicos (CECs) Institutional Animal Care and Use Committee and are in accordance with relevant guidelines and regulations. Universidad Austral de Chile (UACh) investigation animal use bioethics subcommittee approved all experimental procedures.

### Acute lung injury model

Animals were anesthetised with 10mg/kg of xylazyne and 100mg/kg of ketamine intraperitoneal (IP), and nasally instilled three times with lipopolysaccharide (LPS) from *Escherichia coli* in the prophylactic (0.33μg LPS/10μl PBS every 5 minutes) and from *Pseudomonas aeruginosa* in the therapeutic model (0.33μg LPS/10μl PBS every 5 minutes). The control groups of mice were nasally instilled with a total of 30 μl PBS. Animals were sacrificed 24 hr after instillation to obtain BALF. Animals were sacrificed by anaesthesia over-dosage injecting 20mg/kg of xylazyne and 200mg/kg of ketamine I.P. delivered.

### Prophylactic model

TAMX was resuspended in corn oil and administered intraperitoneally at 1mg/200µl. In order to explore the effect of TAMX over inflammatory cells arrival to the airways, mice were treated with the drug or the same volume of corn oil as vehicle (VEH) at days −3, −2, and −1 before LPS instillation. Animals of both groups were subsequently divided resulting the following groups: 1st was injected with the vehicle and instilled with PBS, 2nd injected with TAMX and also instilled with PBS, 3rd injected with the vehicle and instilled with PBS, and the 4th group injected with TAMX and instilled with LPS. Cells in BALF were placed in slides using cytospin and stained with May-Grünwald/Giemsa for the differential count.

### Therapeutic model

After 8 and 24 hr of LPS or PBS instillation, TAMX (1 mg) or same volume of corn oil (vehicle) were injected. The PBS group was divided in two sub-groups: 1st was injected with TAMX while the 2nd correspond to untreated controls. The LPS group was divided in three sub-groups: 1st TAMX injected, 2nd vehicle injected and the 3rd was left as untreated. This design corresponds to the acute lung injury model and BALF samples were obtained 48 hr after instillation. For the study in the chronic disease model, wild type and *Scnn1b*^tg/+^ animals were injected with a single dose of TAM (1 mg), vehicle or left untreated. Animals were sacrificed 24 hr post-injection, to obtain BALF. Later, total cells in BALF were counted in a neubauer chamber, placed in slides using cytospin and stained with May-Grünwald/Giemsa for the differential count.

### Ussing chamber experiments

Tracheae were placed in P2306B of 0.057 cm^2^ tissue holders and placed in Ussing chambers (Physiologic Instruments Inc., San Diego, CA, USA). Tissues were bathed with bicarbonate-buffered solution (pH 7.4) of the following composition (in mM): 120 NaCl, 25 NaHCO_3_, 3.3 KH_2_PO_4_, 0.8 K_2_HPO_4_, 1.2 MgCl_2_, 1.2 CaCl_2_ and 10 D-Glucose, gassed with 5% CO_2_– 95% O_2_ and kept at 37°C. The transepithelial potential difference referred to the serosal side was measured using a VCC MC2 amplifier (Physiologic Instruments Inc.). The short-circuit currents (I_SC_)were calculated using the Ohm’s law as previously described [12]. Differences (Δ I_SC_), were calculated from I_SC_ after minus before the addition of drugs. The experiments were performed in the presence of 10 µM amiloride in the apical side to block sodium absorptive currents. The 1 µM forskolin (FSK) + 100 µM 3-isobutyl-1-methylxanthine (IBMX) cocktail was used to stimulate cAMP-dependent anion secretion, and 100 µM uridine-5′-triphosphate (UTP) to evoke Ca^2+^-activated anion secretion.

### Statistical analysis

SigmaPlot 12.3 was used for graph generation and statistical analysis. Shapiro Wilks was used to test the distribution of the data. Kruskal Wallis test was performed to compare the differences between groups in each assay, and this was followed by a Tukey or ANOVA on ranks tests. Overall, p values <0.05 were considered as significant.

## RESULTS

### Tamoxifen reduces airway neutrophilia in mouse models of acute lung injury and chronic muco-obstructive lung disease

To test if tamoxifen can be used as a therapeutic drug, we used a mouse model of acute lung injury induced by LPS instillation either through a prophylactic or a therapeutic approach. Administration of TAMX previous to the induction of the lung injury results in a decreased percentage of neutrophils in the airway (Figure 1). Following a similar approach, we explored the use of a double dose of TAMX (8 and 24h post LPS instillation) as treatment. Analysis of cellularity demonstrated that LPS induced an increase in total cells (Fig 2A) that is exclusively due to neutrophils (Fig 2B), as macrophages (Fig 2C) and lymphocytes (Fig 2D) did not changein quantity. As observed in Fig 2B TAMX treatment decreased neutrophil infiltration, impacting total cell number accordingly (Fig 2A).

**FIGURE 1.**
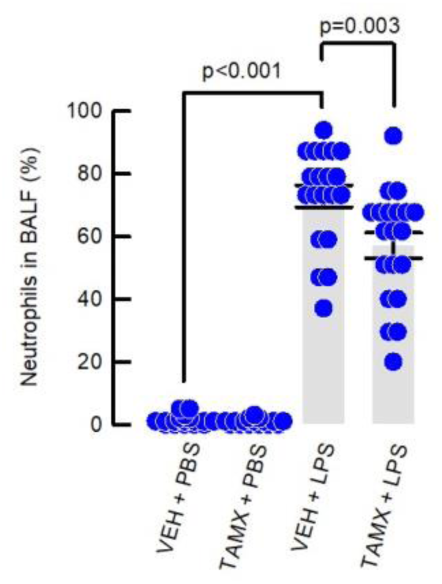
Prophylactic administration of TAMX decrease neutrophils infiltration of the airways in ALI challenge. Percentage of neutrophil infiltration in the airways of mice instilled with LPS or PBS, in animals previously injected with TAMX or VEH once a day for tree previous days before LPS instillation (n=18 in VEH+PBS and TAMX+PBS; n=20 in VEH+LPS and TAMX+LPS). Differences were calculated (p) using ANOVA on ranks.

**FIGURE 2.**
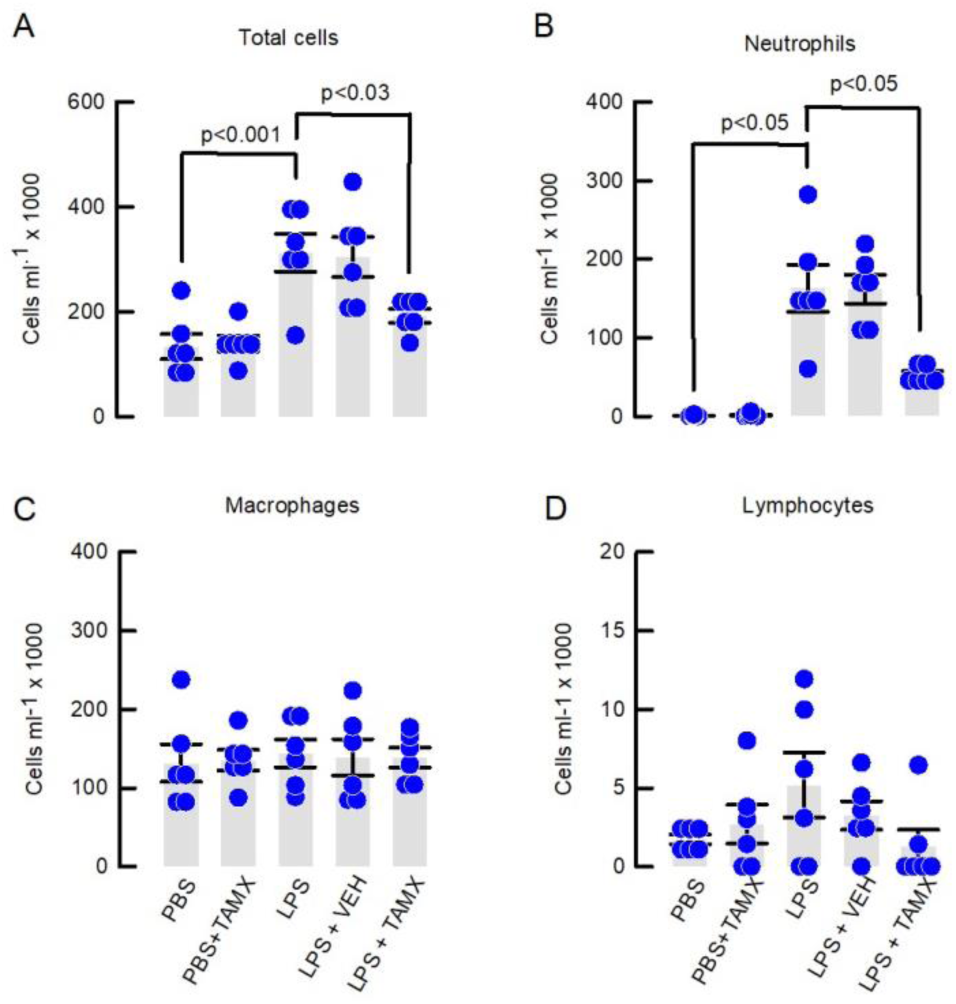
Therapeutic administration of tamoxifen decrease neutrophils infiltration of the airways in acute lung injury. Leukogram summarizing the total cells (A), neutrophils (B), macrophages (C) and lymphocytes (D), obtained from BALF of wild type animals instilled with LPS and treated with TAMX at 8 and 24h after LPS instillation. Differences were calculated (p) using ANOVA on ranks.

We extended our results testing TAMX in a model of chronic muco-obstructive disease, the *Scnn1b*^tg/+^ mouse. This animal is affected by chronic airway inflammation with infiltration of inflammatory cells and release of pro-inflamamtory cytokines characteristics of CF and COPD [13]. We used a single dose of TAMX and obtained BALF samples 24h later. As observed in Figure 3A the *Scnn1b*^tg/+^ airways showed increased number of total inflammatory cells when compared to controls and TAMX treatment decreased the number of total inflammatory cells in BALF. Analysis of cell types demonstrated that the neutrophilia was decreased after TAMX treatment in the *Scnn1b*^tg/+^ mice. No effect of TAMX was observed in macrophages (Fig 3C) or lymphocytes (Fig 3D) numbers.

**FIGURE 3.**
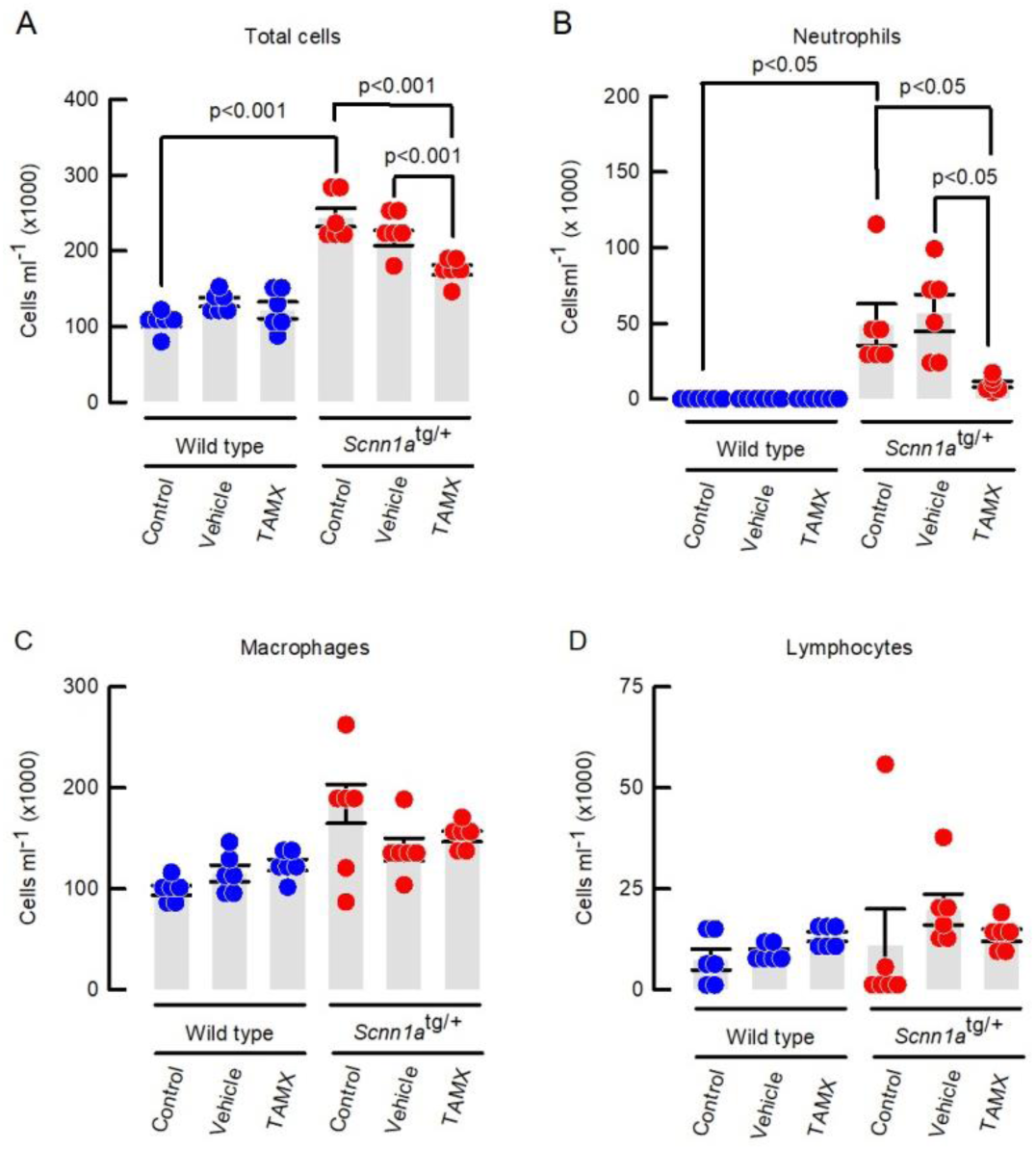
Tamoxifen reduced neutrophil infiltration in the *Scnn1b*^tg/+^ mouse airways. Leukogram summarizing total cells (A), neutrophils (B), macrophages (C) and lymphocytes (D) obtained from BALF of wild type and *Scnn1a*^tg/+^ animals treated with TAMX. p values were calculated using ANOVA on Ranks. Blue dotes for control animals (wild types) and red dots for *Scnn1b*^tg/+^ mice.

### Tamoxifen does not affect electrophysiological parameters in mouse airways

TAMX has been reported to block several ion channels including chloride [14,15]. Absence of anion secretion, as observed in CF, is absolutely detrimental for airway function, impairs airway surface hydration, negatively affect mucus deployment, and triggers a devastating inflammatory response [16]. Thus we tested if TAMX affected anion secretion in the trachea of mice. As observed in Fig 4A, the addition of tamoxifen (0.1-10 µM) on the apical side of the epithelium does not affect basal currents, and subsequent addition of amiloride to unmask Na^+^-absorption, IBMX+forskolin to activate cAMP-dependent anion secretion and UTP to induce Ca^2+^-activated anion secretion was unaltered, as the values obtained where not different from those calculated from control tissues or those treated with DMSO (Fig 4B).

**FIGURE 4.**
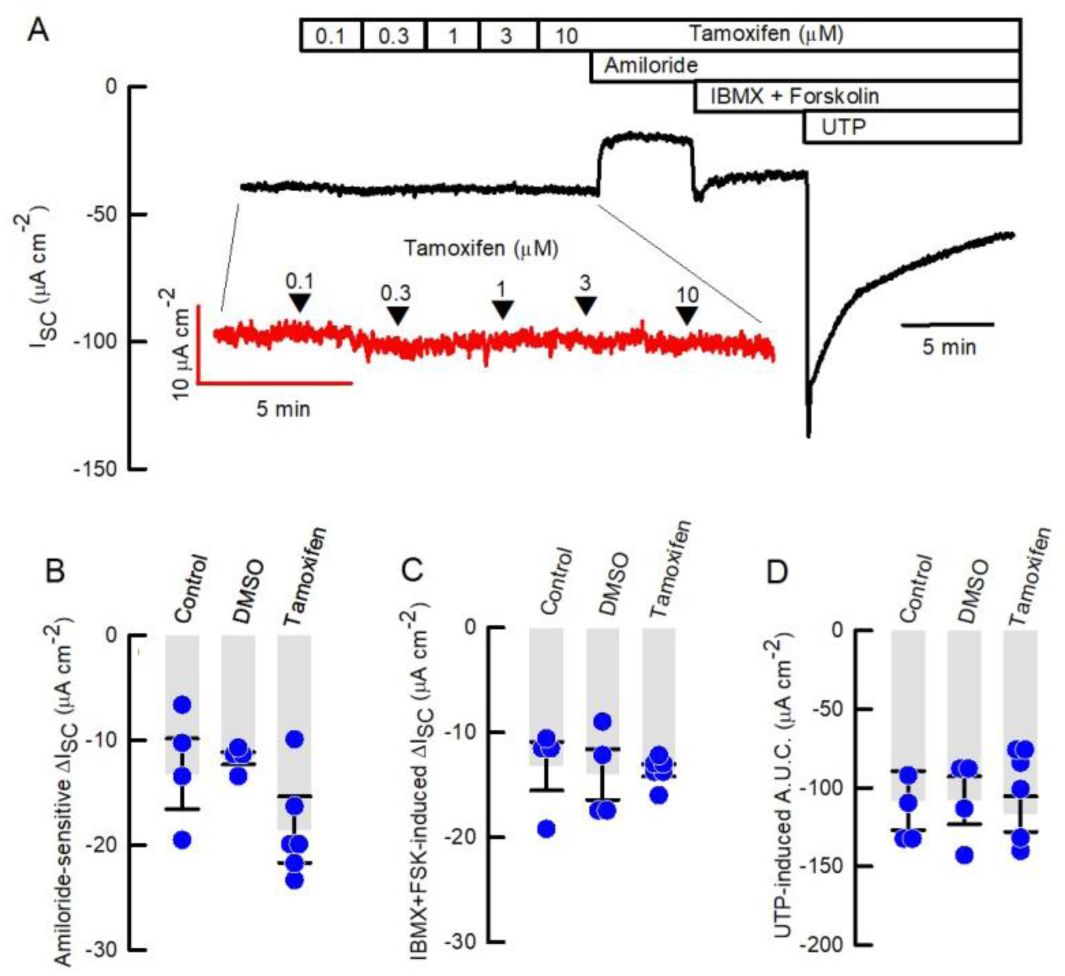
Tamoxifen does not affect ion absorption and secretion in the mouse airways. (A) representative short-circuit current recording from mouse trachea to study the effect of apical TAMX (0.1-10 µM) on basal current (amplified in the red trace), Na^+^-absorption (amiloride 100 µM), cAMP-induced anion secretion (IBMX 100 µM +Forskolin 1 µM) and Ca^2+^-activated anion secretion (UTP 100 µM). Summary of amiloride-sensitive (B), cAMP-induced (C) and Ca^2+^-activated anion secretion (D), for control, DMSO and TAMX. No statistical differences were found (p>0.05); ANOVA on Ranks.

## DISCUSSION

Neutrophil recruitment must be in equilibrium with their removal from inflamed tissues or undesired damage and often death is the result for many human patients. Due to the current worldwide increase in inflammatory diseases affecting the lungs, there is urgency for new and better treatments. We studied TAMX, a drug of choice for the treatment of breast cancer, and observed a significant reduction of neutrophil infiltration of the airways using three animal models: prophylacltic and treatment for ALI, and in a transgenic mouse affected by chronic inflammation of the airways. Our results demonstrated an important decrease in the amount of infiltrated neutrophils using TAMX in all cases.

The effects of TAMX in airway disease have been previously observed by us and others. In horses affected by EAS and after acute exposure to antigens, TAMX treatment produced a decrease in the amount of BALF neutrophils and improvements of the clinical status of the animals [9]. Nevertheless, during a chronic-exposure to antigens TAMX produced an important reduction in airway resistance but failed to reduce neutrophil infiltration [17]. The mechanism of neutrophil reduction seems to be due to the pro-apoptotic effect of TAMX [18], that induces phosphatidylserine exposure in the outer cell-membrane of neutrophils, facilitating it’s removal from the inflammatory *milieu* by promoting efferocytosis and therefore the resolution of inflammation [19].

The current results reproduced our previous findings in horses, now using three different approaches in mice. Animals that received TAMX previous to airway instillation of LPS showed reduced amounts of neutrophils collected in BALF. The same results were observed when TAMX was administered after LPS instillation, demonstrating that TAMX can reduce the amount of neutrophils arriving to the lungs in response to acute lung injury. Interestingly, a single dose of TAMX reduced the number of neutrophils in the airways of the CF/COPD mouse model *Scnn1b*^tg/+^ whose inflammatory state arose from the dehydration of the airway surface fluid layer (ASL). In a similar fashion, a different COPD model induced by smoke-inhalation showed to be protective against increased TGF-beta and superoxide production in the lungs, when receiving TAMX for 2 weeks [20]. The fact that TAMX can act as a blocker of ion channels is a possible drawback of its use in inflammatory airway disease, but the results observed in the *Scnn1b*^tg/+^ animals reduced this chance. Moreover, TAMX has been proved to have a pro-secretory effect in human airway epithelium that might cooperate in the reduction of the inflammatory response of the airways by restoring ASL volume and mucociliary functions, discarding again, any inhibitory effect on anion secretion [21]. Finally, our Ussing chamber experiments demonstrated that the addition of TAMX do not affect basal electrical parameters or absorptive and secretory currents in the airway epithelium of the mouse trachea, supporting the idea that TAMX does not affect ASL-volume in a negative fashion.

Previous information indicated that among other NSAIDs, TAMX was more effective at suppressing NF-kB in tumour cells than dexamethasone [10]. The glucocorticoids (including dexamethasone) are commonly used as a therapeutic tool for some phenotypes of asthma [22]. However, patients with severe neutrophilic asthma are often refractory to treatments with glucocorticoids as they affect neutrophils [23], enhancing their arrival by upregulating the epithelial expression of IL-8 [24] and increasing neutrophil’s survival [25–27]. A similar result was obtained in *Scnn1b*^tg/+^ animals treated with prednisolone (another glucocorticoid), that did not prevent neutrophil infiltration of the airways [28].

In summary, we conclude that TAMX is effective at reducing neutrophil infiltration of the airways of mice subjected to ALI by LPS instillation and in a model of non-infectious chronic airway inflammation. As the side effects of TAMX treatment are long-term [29], TAMX might pose an effective shot-term treatment to reduce neutrophil infiltration of the airways.

## ACKNOWLEDGEMENTS

CECs is funded by the Centers of Excellence Base Financing Program of CONICYT. Supported by FONDECYT 11160418 (C.H.).

